# A Web Service for Automated Design of Multiple Types of Ribozymes Targeting RNA: from minimal hammerhead to aptazymes

**DOI:** 10.1101/2023.09.30.560155

**Authors:** Sabrine Najeh, Nawwaf Kharma, Thomas Vaudry-Read, Anita Haurie, Christopher Paslawski, Daniel Adams, Steve Ferreira, Jonathan Perreault

## Abstract

Ribosoft 2.0 is the second version of a web service to design different types of *trans*-acting conventional and allosteric ribozymes. The web service is publicly available at https://ribosoft2.fungalgenomics.ca/. Ribosoft 2.0 uses template secondary structures that can be submitted by users to design ribozymes in accordance with parameters provided by the user. The generated designs specifically target a transcript (or, generally, an RNA sequence) given by the user. Herein, sixty ribozymes of different types were tested on two different mRNAs, with a majority shown to be active. We have also generated and proved the activity of the first *trans*-acting aptazyme designed *in silico*.

## INTRODUCTION

Ribozymes are non-coding RNA with catalytic activities. Ribozymes were discovered in 1981 in a study demonstrating that the effector of the splicing mechanism was an RNA molecule rather than a protein, and those RNAs were designated group I introns (1,2). This event led to the discovery of many other ribozymes, including a group of short RNAs sharing a common ‘transesterification’ reaction. This type of reaction occurs at a specific site, causing the self-cleavage of the RNA through a nucleophilic attack, generating an RNA molecule with a 2’-3’ cyclic phosphate and a 5’ hydroxyl group. RNAs belonging to this group are called ‘small self-cleaving RNAs’, and they are classified into different families based on the secondary and tertiary structure that they adopt, in order to affect the cleavage reaction (3). The hammerhead ribozyme (HHR) was the first small self-cleaving ribozyme to be discovered and is the one most studied. HHR was first discovered in plant-infecting viroids, before subsequently being discovered in all domains of life, including prokaryotes and eukaryotes, ranging from single-cell organisms to mammals including humans (4). Since the discovery of HHR, many different families of short self-cleaving RNAs have been identified, including hepatitis delta virus (5), GlmS (6), hairpin (7), Varkud satellite (8), twister (9), twister sister, hatchet and pistol (10) ribozymes. Some of these ribozymes are involved in viral and viroid replication cycle (11,12), while the natural function of others remains less-well defined. Ribozymes were also used for gene expression control purposes by their inclusion into mRNA sequences, downstream or upstream of coding sequences (13). They have also been used as independent *trans-* cleaving RNA molecules that target mRNAs anywhere, including in their coding sequences (13,14).

Self-cleaving ribozymes possess many potential advantages for RNA targeting tool development and gene expression control. In general, this advantage is due to their small size and activity which is often independent of protein helpers, making them easier to test *in vitro*. Besides their ability to self-cleave (*cis*-cleavage), ribozymes can be conceptually divided into two parts: a ribozyme strand and a substrate strand, with no loss in activity. The binding site of the ribozyme strand can be easily modified to recognize and hence, cleave other RNA targets, at specific sites (*trans*-cleavage) (15).

The majority of small self-cleaving RNAs do not need cofactors for their activity and they are active by default. This latter characteristic can be a limiting factor, when such ribozymes are used in environments where their activity must be controlled. To respond to this need, allosteric ribozymes were developed (16). One method used to develop ribozymes with allosteric (modulator-dependent) activity is to link the ribozyme sequence to an aptamer, which is known to bind efficiently and specifically to a desired target (17–19). In this context, an aptamer will interfere and disturb ribozyme folding to disrupt its activity. Binding of the cofactor to the aptamer sequence will alleviate this interference and subsequently rescue ribozyme activity. The combination of a ribozyme and an aptamer is often called ‘aptazyme’. Generally speaking, aptazyme activity depends on the presence or absence of the cognate molecule of the aptamer, where the structure of the aptamer bound to its ligand can either disrupt or stabilize the active structure of the ribozyme module. Aptazymes were used in different applications, such as the control of gene expression and viral vector replication (18,20–23)

HHR has a conserved structure of three stem-loops with a catalytic core of 17 nucleotides specific to the cleavage site GUH (Figure 1). To generate a *trans*-acting version of a HHR, stem II and the catalytic core must be conserved, stem I and III are modified to be perfectly complementary to the target substrate (24). Nevertheless, making a new design cannot be achieved by simply changing the ribozyme sequence; other parameters should be taken into consideration. The ability of a modified sequence to fold into the hammerhead ribozyme’s active structure is an essential condition of catalytic activity. Also, the lengths of stems I and III must offer sufficient target specificity, while simultaneously allowing for some ribozyme turnover. Making stems I and III too long will inhibit turnover and excessively short stems raises the risk of reduced activity and increased off-targets effects. Hence, even if manual designs can often be functional, they are much less likely to be optimal. The manual process is also time-consuming and offers limited design choices. A bioinformatics tool taking into account all the important parameters to automatically generate designs would greatly improve this process, reduce human effort (and error), as well as open the door to many ribozyme and aptazyme applications.

**Figure 1:**
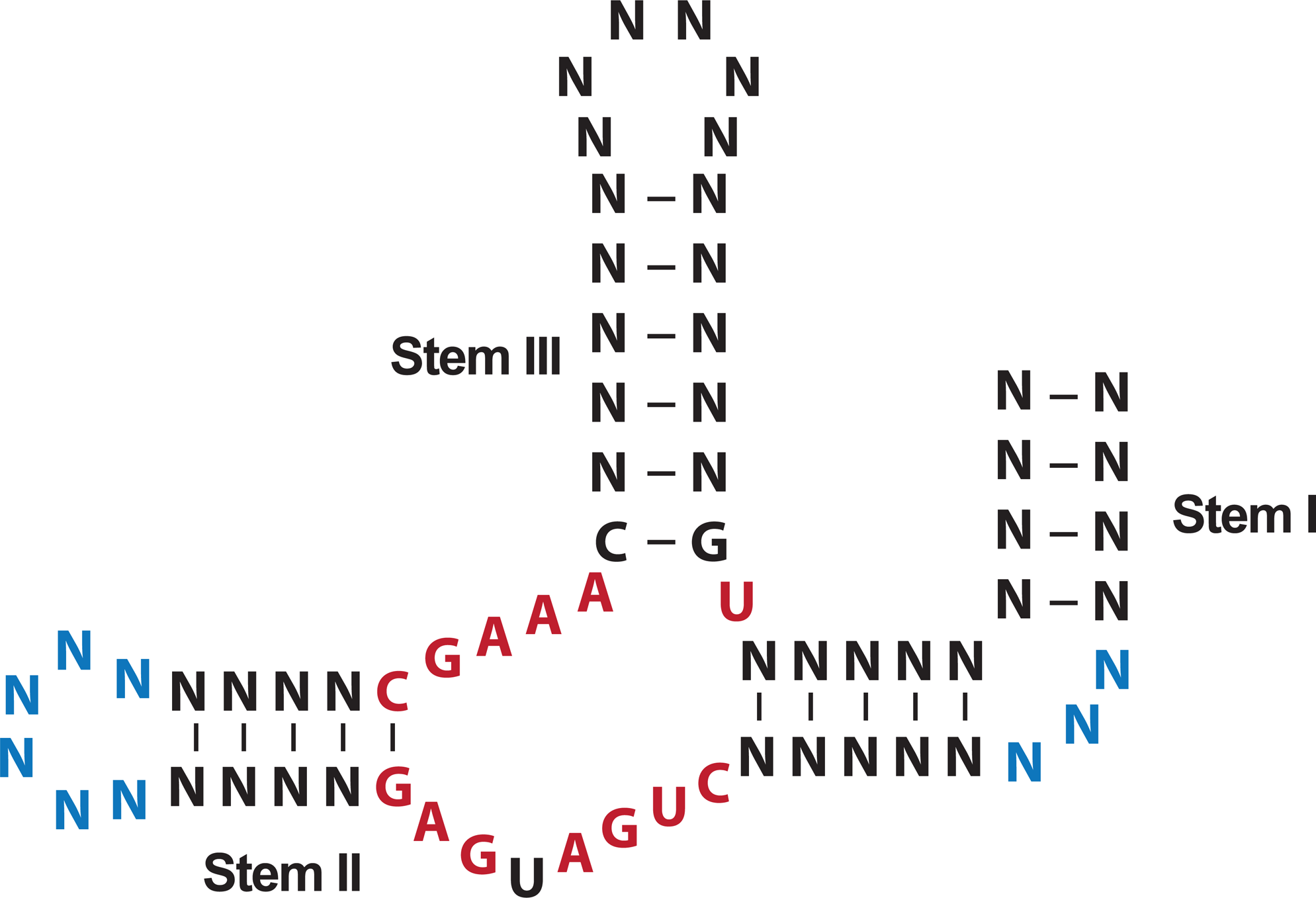
Hammerhead ribozyme structure. Three stem loops surround the conserved catalytic core (red nucleotides). The blue sequences are nucleotides that form a tertiary interaction (a pseudoknot for example) that occurs between stem I and stem II.

In this paper, we present Ribosoft 2.0, an improved version of Ribosoft (25) for the design of *trans*-acting ribozymes. This new version of the web service offers the ability to design different conventional and allosteric ribozymes, with known secondary structures. As opposed to similar webservers, designs can be made for almost any known ribozyme. Different types of ribozymes can be designed from the web service: different types of HHR (extended or pseudoknotted version), pistol, twister, twister sister (26,27) and aptazymes controlled by FMN (28). This program is able to generate sequences of active ribozymes against any desired target RNA. During the process of generating new sequences, Ribosoft 2.0 evaluates different parameters. The ribozymes structure is validated by RNA fold (29) to make sure the new sequence folds into the same structure as the template. The accessibility of the cleavage site is calculated using the target RNA’s secondary structure. Finally, the specificity of the ribozyme sequence (off-targets effects) is evaluated by BLASTing the target sequence on NCBI (30). We used Ribosoft 2.0 to design ribozymes of different types, to target mRNAs encoding Bcl2 and Galectin7 proteins as example targets.

## MATERIALS AND METHODS

### Software Methodology

The *backend* of the Ribosoft 2.0 web service relates to the correct processing, via programmed algorithms, of the biological information provided by the end-user or accessed by the program.

Known as an *activity diagram*, Figure 2 provides an overview of the whole program, with each vertical section representing a well-defined functionality. The *front end,* which is best presented as a Graphical User Interface (or GUI) deals with end-user interaction: input from the user (Figure 3) and ranked design output (Figure 4). The core information processing modules of Ribosoft 2.0 are all part of the back-end and the algorithm library, each component of these two sections (e.g., generate design or evaluate accessibility) are described, using natural language and pseudo-code below.

**Figure 2:**
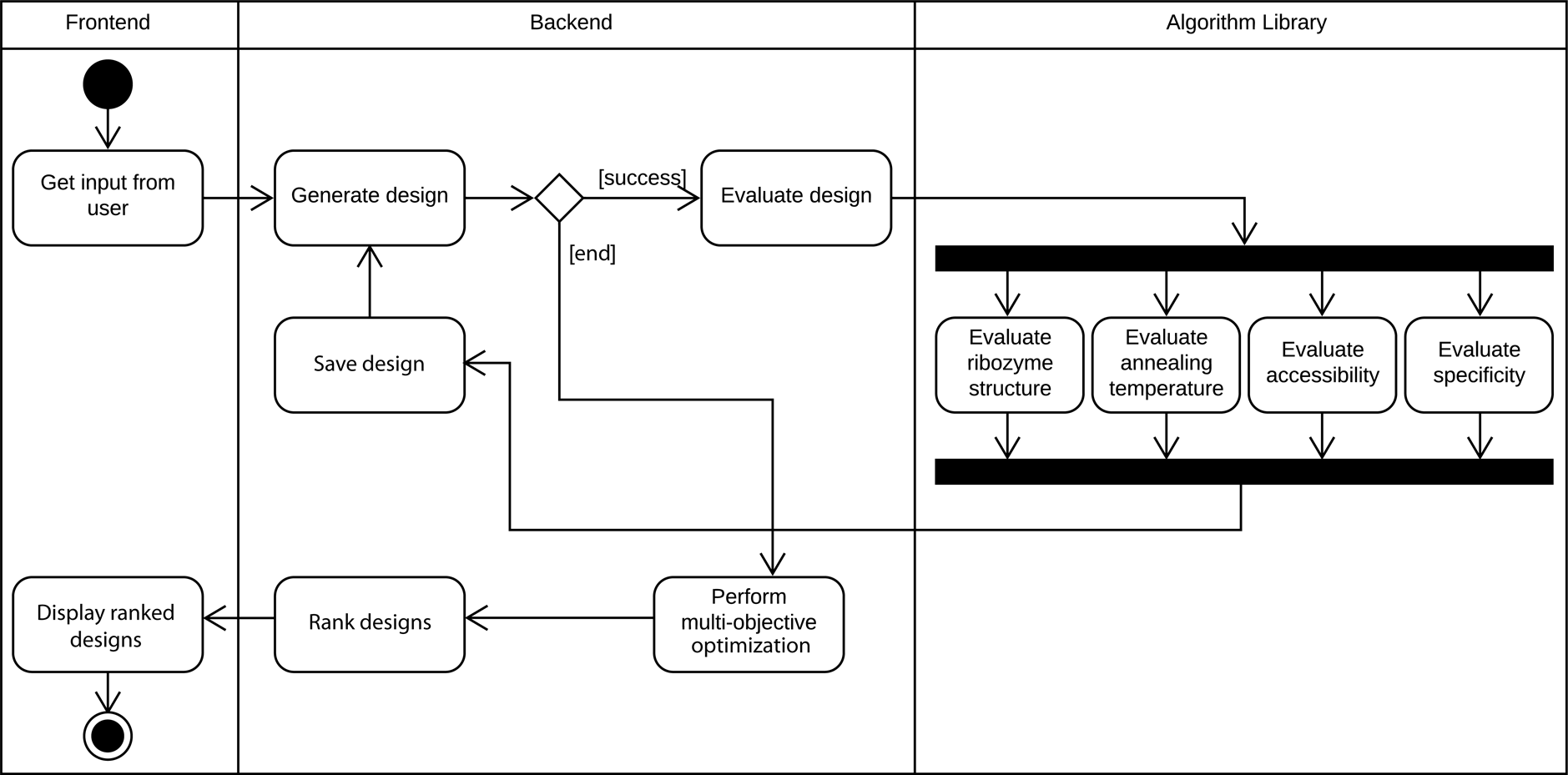
Complete overview of Ribosoft (V2) program activity.

**Figure 3:**
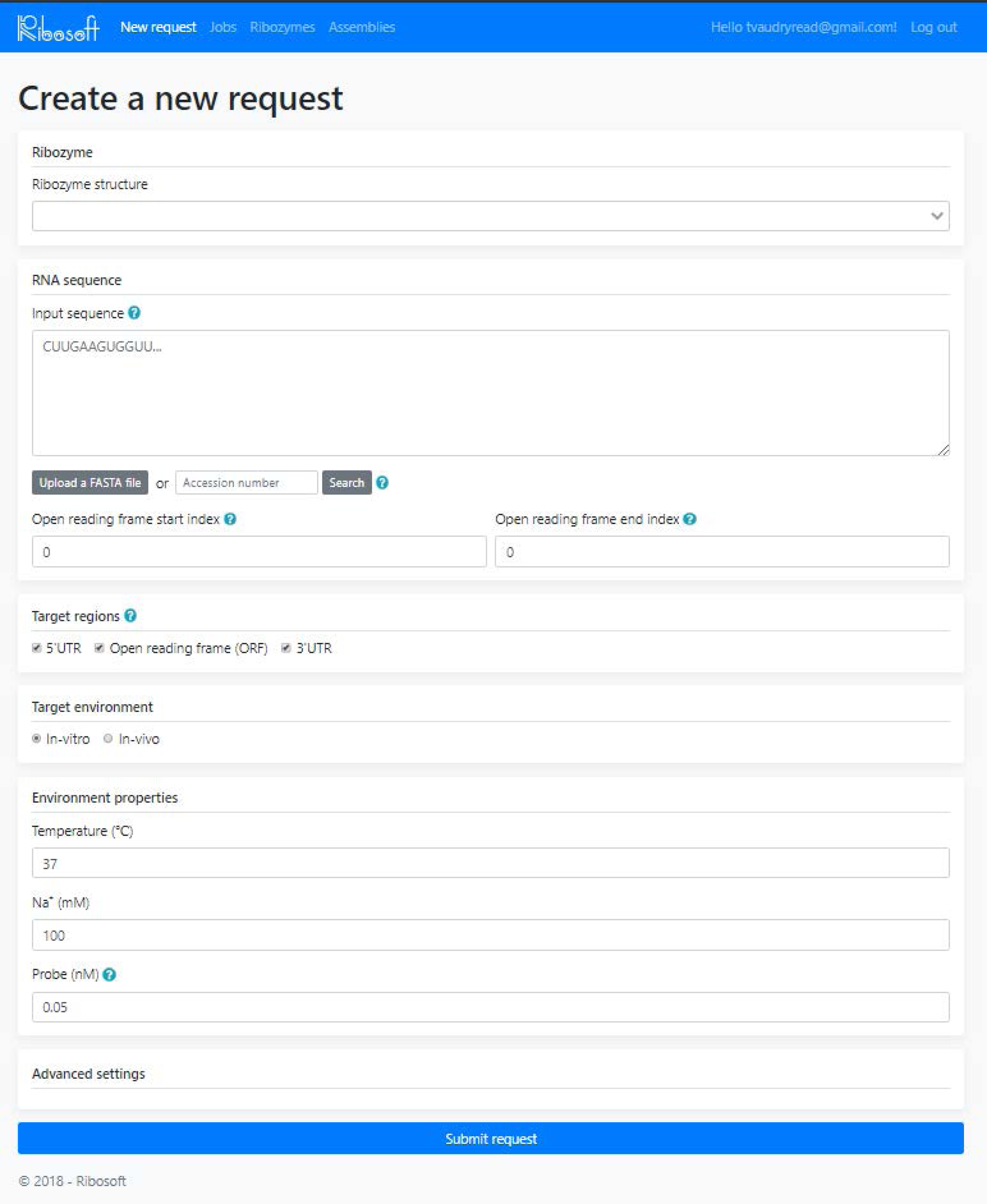
Input specifications provided by user.

**Figure 4:**
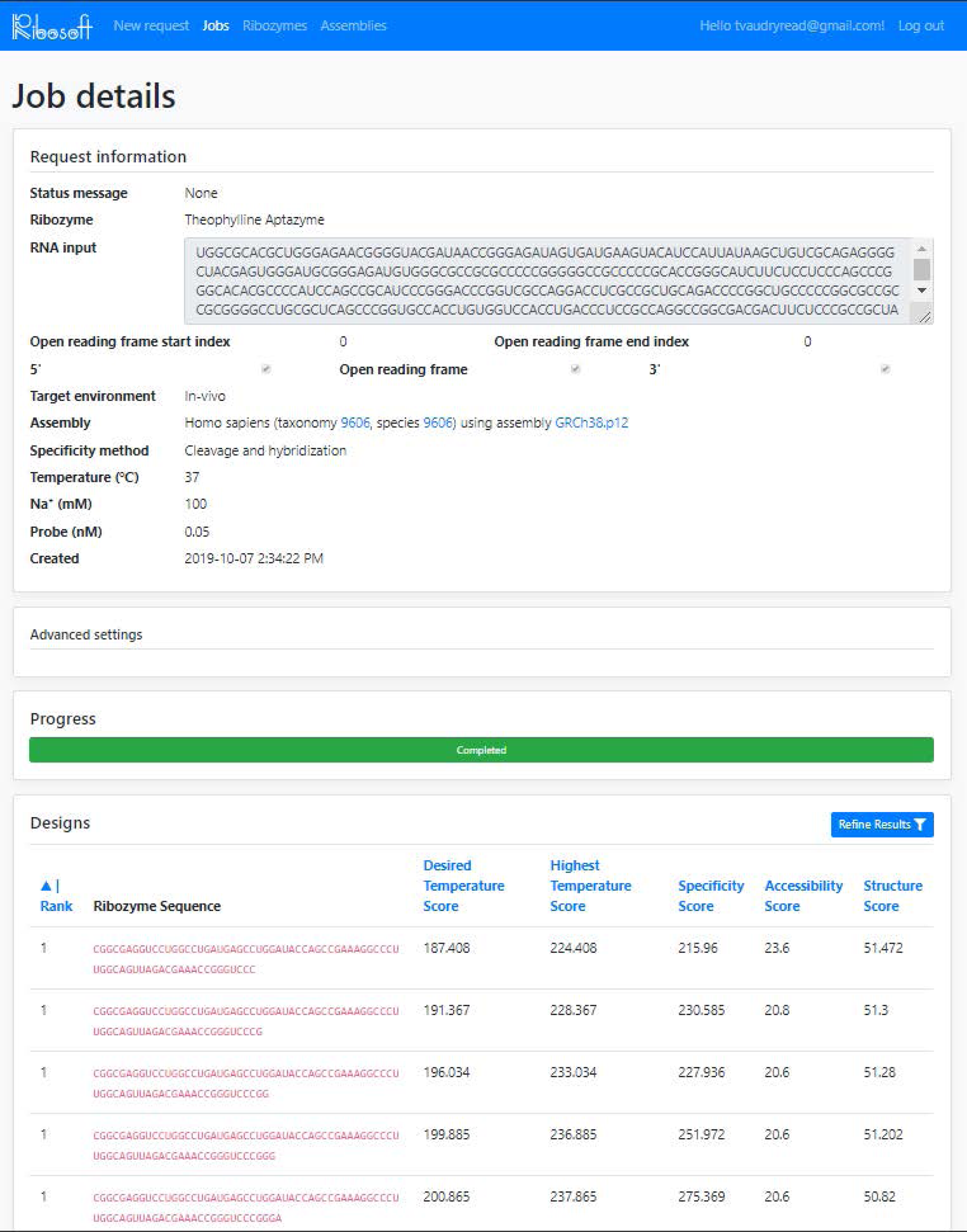
Output designs presented to user.

### Back-end and Algorithm Library

#### A. Design Generation

Every instance (specific design) of a general template representing the used ribozyme is generated. Hence, the program discards all ribozyme instances that do not hybridize (strongly enough) to the binding regions around legitimate cleavage-sites on the provided transcript (of the targeted gene/sequence). The instances that remain are then evaluated for four different fitness terms: ribozyme structure, annealing temperature, cut-site accessibility and specificity (to the target transcript and not others within the chosen organism). Hence, the program stores all evaluated designs with their fitness terms in a database.

#### B. Design Objectives evaluation

##### B1. Evaluate Ribozyme Structure

~~~
Fold ribozyme sequence // this generates multiple foldings
*Score* = 0
**FOR** each *folding*
    Calculate distance between *folding* and *ideal folding
    Score* = *Score* + *distance \** probability of *folding*
**END FOR
RETURN** *Score*
~~~

##### B2. Evaluate Annealing Temperature

~~~
Determine *limits* of binding regions
*Score* = 0
**FOR** each *binding region*
    Calculate *annealing temperature* of binding region
    *Score* = *Score* + *annealing temperature*
**END FOR
RETURN** *Score*
~~~

##### B3. Evaluate Accessibility

~~~
Fold whole transcript, without constraints
Calculate the minimum free energy of the resulting folding (MFE)
Determine the limits of the binding regions on the transcript
Fold whole transcript with constraints (forcing open all binding regions)
Calculate the minimum free energy of the constrained folding (MFE_R)
*Delta* = MFE_R – MFE
**RETURN** *Delta*
~~~

##### B4. Evaluate Specificity

~~~
Form a search string using the binding regions of the ribozyme
**BLAST** the search string for chosen organism to:
     (1) find the *number* of affected transcripts
     (2) **FOR** *each* transcript:
         find the *percentage* of overlap with the search string
Calculate a *Score* reflecting (1) and (2)
**RETURN** *Score*
~~~

#### C. Multi-Objective Optimization & Ranking

Multi-objective optimization is the last step in the design process, receiving input from the job and outputs a ranked list of candidates for the user. It is important that this step be quick and accurate, as it is the final step to complete a job. The advised method for ranking is Pareto Dominance, for its simplicity and generality.

### Generation and selection of ribozymes for experimental assays

Before designing ribozyme sequences, different types of ribozymes were added to Ribosoft 2.0 and our now available in the dropdown list. This was done by providing models such as that of the pistol ribozyme (Figure 5A) as well as four different hammerhead ribozymes, including aptazymes (Figure 5 and supplementary material). Afterwards, we used Ribosoft 2.0 as any user should by simply adding our sequence as shown in Figure 3. Parameters for environment properties that we used as input were: 37°C, 100 mM NaCl and 0.05 nM probe, with 0.05 for advanced settings of all parameters determined by Ribosoft 2.0, this number relates to a chosen range to consider values equivalent with regards to the Pareto Dominance.

**Figure 5:**
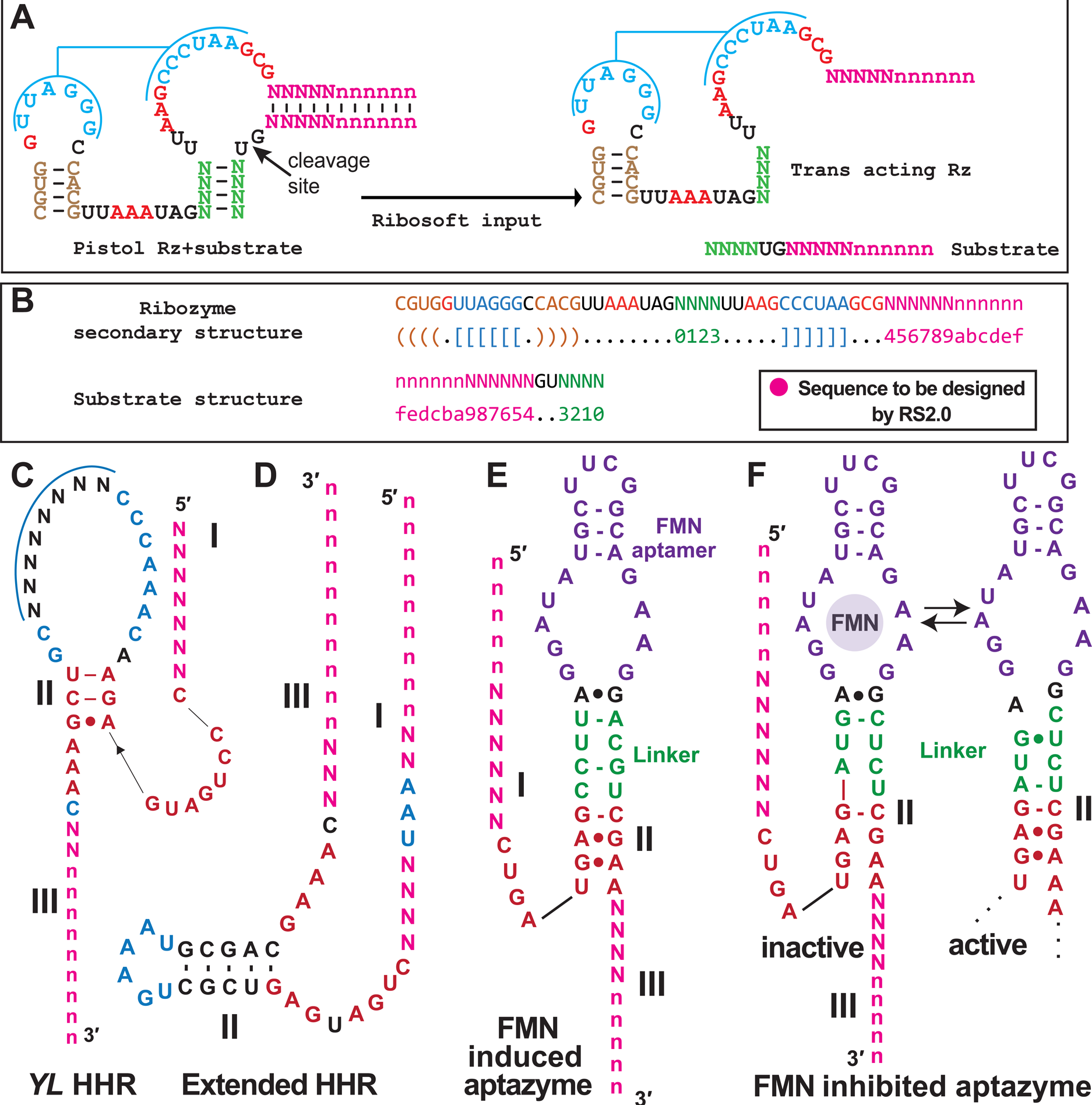
*Trans-*acting ribozymes designed by Ribosoft 2.0. (A) Input of new ribozyme models for Ribosoft2.0. Left: Pistol ribozyme structure is formed of three stems and two loops. In red is the conserved sequence of the catalytic core. Green and pink nucleotides are the sequences to be modified by Ribosoft depending on the target sequence. Blue color shows the sequences that form a pseudoknot. Right: Input of Ribosoft 2.0 : Pistol ribozyme secondary structure divided in a ribozyme strand (with the catalytic core) and a substrate strand containing the cleavage site. (B) Each strand structure is given to Ribosoft 2.0 as input where ‘.’ is a single stranded part, ‘( )’ is a base pair and ‘[ ]’ is a pseudoknotted part. To figure out the complementarity between ribozyme and substrate strands, the sequences letters and numbers are used in the same order from both sides. (C), (D), (E) and (F) Secondary structure of the ribozyme strands. Templates used by Ribosoft 2.0 to design new sequences (C) HHR *YL*, (D) extended HHR, (E) FMN-induced aptazyme and (F) FMN-inhibited aptazyme, with both inactive (FMN-bound, according to (50), used by Ribosoft to design sequences) and active structures pictured. For all the ribozymes the catalytic core (red nucleotides) and one of the stems was conserved (stem II for all the HHR and P1 for the pistol ribozyme). The nucleotides in pink ‘Ns’ are the single stranded stems that will be designed by RS 2.0 to be complementary to the target sequence surrounding the cleavage site of each ribozyme.

Sixty ribozyme sequences designed by Ribosoft 2.0 were chosen to be assayed. All the ribozyme sequences were chosen to pair the highest and lowest scores for the following Ribosoft 2.0 parameters: 1- “temperature”, 2- “accessibility” and 3- “structure”; “specificity” was not considered as *in vitro* characterization of ribozymes would not provide any relevant data in that regard. For these ribozymes, 30 were designed to target Gal7 mRNA and 30 to target Bcl2 mRNA. (Supplementary Table S1).

### Preparation of transcription template

DNA templates used for the ribozyme production were purchased as ssDNA oligos (Alpha DNA or IDT), double stranded DNA templates were produced by primer extension. Oligonucleotides were incubated with a PCR mix containing (100 µM dNTP, 1X thermopol Taq reaction buffer, 1X of Q solution (Qiagen) and 1 µM of each primer). A complete list of oligonucleotide sequences is available in supplementary Table S1.

The expression vector (Pet22b G7h) was used as a template for the amplification of the Galectin7 gene by PCR. The forward primer used harbors the T7 promoter. dsDNA product, was purified and used as template for *in vitro* transcription, using T7 RNA polymerase, to generate Galectin7 mRNA.

### Reverse Transcription-PCR

Total RNA was extracted from Jurkat cells (10^6^ cells) using Trizol then treated with DNase I to eliminate the genomic DNA. Bcl2 cDNA was generated using the one-step RT-PCR Qiagen Kit containing 1U of Enzymes mix (Omniscript and Sensiscript Reverse Transcriptases and HOTstart Taq DNA polymerase, provided one-step RT-PCR buffer (5x) Q solution (5x), 2 mM dNTP mix and complete to 50 µl with RNase free water. Primers used were designed to amplify a specific segment of the Bcl2 cDNA.

### Transcription

The ribozymes synthesis was made by *in vitro* transcription in presence of the PCR product for each ribozyme. Each PCR product was incubated with 3 mM rNTPS (2 mM each of rATP, rGTP, rCTP and rUTP), 1X transcription buffer (80 mM HEPES-KOH pH 7.5, 24 mM MgCl_2,_ 40 mM DTT, 2 mM spermidine), 1 U/μl of inorganic pyrophosphatase Roche), USA, 1 U/μl RNase inhibitor (Thermofisher), 1 U/μl T7 RNA polymerase and Milli-Q water for 2 hours at 37°C.

After reaction completion, 1 U/μl of DNase I (RNase free) (NEB) was added to each reaction followed by 30 min incubation at 37°C for the elimination of the DNA templates. RNA products were extracted with phenol/chloroform, then precipitated with ethanol overnight at −20°C. They were, afterwards, fractioned on a 10% denaturing polyacrylamide gel. The band corresponding to the expected size of each ribozyme was cut and the RNAs were extracted overnight in the elution buffer (EDTA 0.1 mM, NaCl 0.3 M, SDS 0.001%) at 4°C. Finally, the RNAs were precipitated by ethanol, dried, resuspended in water and their concentration measured using a nanodrop spectrophotometer.

To be able to visualize the cleavage activity of the ribozymes, the target RNAs were either externally (5’) radiolabeled, when this was not possible, due to target RNA length and structure, we used internally radiolabeled RNA targets made by radioactive transcription.

### Radioactive transcription

The radioactive transcription for target RNAs (Gal7, Bcl2) was done using the same procedure for the cold transcription except for the use of 0.5 µl alpha-^32^P-labeled UTP (3000 Ci/mmol-10 mCi/ml) (Perkin Elmer) and 0.8 mM UTP (instead of 2 mM). After the DNase I treatment, the RNAs were precipitated with ethanol and purified using a 6% denaturing polyacrylamide gel.

### RNA 5′ labeling

Unlabeled purified RNA targets were dephosphorylated using 1 U antarctic phosphatase (NEB) incubated with 1X of provided buffer during 20 min at 37°C and then 5 min at 65°C to inactivate the enzyme. For labeling, RNA was incubated with 1 U T4 polynucleotide kinase (NEB) in presence of its buffer (70 mM tris-HCl pH 7.6, 10 mM MgCl_2_, 5 mM DTT) and γ-^32^P-ATP. The reaction was incubated for 1 h at 37°C. The 5′-radiolabeled RNA targets were purified on a 6% denaturing polyacrylamide gel, eluted overnight at 4°C and then precipitated.

### End point cleavage assays

Each ribozyme was pre-incubated in the cleavage buffer (50 mM Tris pH 7.5, 100 mM NaCl and 25 mM KCl) at 85°C for 1 min and then snap-cooled on ice. Afterwards, each corresponding target was added to the reaction as well as MgCl_2_ (10 mM) to start the reaction. For the FMN activated and FMN inhibited aptazymes 200 μM of FMN is also added to the reaction. The incubation lasted one hour at 37°, and was stopped by adding an equal volume of the 2X stop dye (formamide buffer: 95% formamide, 10 mM EDTA, 0.025% bromophenol blue and 0.025% xylene cyanol blue).

The resulting cleaved RNAs were fractioned on a 6% denaturing polyacrylamide gel. Then the gel was exposed to a storage phosphor screen overnight. The phosphor screen was scanned with a Typhoon FLA9500 (GE Health Care) and the cleavage extent was quantified using ImageQuant.

### Cleavage time course

To be able to determine the cleavage efficiency and rate of each ribozyme, we followed the cleavage reaction for 64 minutes and aliquots were taken during this time.

The reaction was prepared in the same way for the end-point cleavage assay, but before the MgCl_2_ is added to the reaction an aliquot for the T0 is taken. Once MgCl_2_ is added in the reaction, the timer is started and aliquots are taken at 30s, 1’, 2’, 4’, 8’, 16’, 32’ and 64’. The reaction of each aliquot is stopped with the addition of the stop dye. The results were visualized on 6% denaturing polyacrylamide gels exposed overnight to a storage phosphor screen. The quantification of band intensity was done using ImageQuant and the cleavage rates (*k*_obs_) were calculated with GraphPad where the analysis was done with the ‘one phase decay’ equation.

## RESULTS

To evaluate efficiency and reliability of our web service for RNA design, we validated experimentally the sequences it generates. Ribosoft 2.0 accepts the secondary structure of different types of ribozymes (conventional and allosteric) as input. From these, we have chosen to design 60 ribozymes of five different types. These are the HHR with a pseudoknot (from *Yarrowia lypolitica* genome), extended HHR (from the negative strand of tobacco ringspot virus (sTRSV) modified by (31)), pistol, as well as FMN activated and FMN inhibited ribozymes. And, to determine the efficiency of Ribosoft 2.0 in designing active ribozymes, we targeted these ribozymes against Gal7 and Bcl2 mRNAs. We also aimed to figure out if there is a correlation between the ribozyme activity and the parameters calculated by Ribosoft 2.0 (for ribozyme structure, accessibility and highest temperature scores). For each parameter, we picked a ribozyme with the highest score and another with the lowest score, in both cases targeting Bcl2 and Gal7 mRNAs (supplementary Table S2), which encode for an anti-apoptotic protein (32,33) and a lectin (34), respectively, both with pro-cancer impact (35,36).

### Comparison of end-point cleavage of pistol, extended HHR and pseudoknotted HHR

For hammerhead and pistol ribozymes, 36 ribozymes were tested. Results show that 15 of 18 ribozymes tested against Galectine7 mRNA were active (Figure 6), with cleavage efficiency ranging from 22 to 93%. In total, close to half of these ribozymes, 7 out of 18, cleaved their designated targets at over 50% efficiency, and similar results were obtained with 18 ribozymes against a 719 bases fragment of Bcl2 mRNA (data not shown). These are relatively good results for ribozymes acting on long mRNA-like targets (37–40), since less active ribozymes often require more than 1 hour to result in cleavage efficiencies exceeding 50% (41).

**Figure 6:**
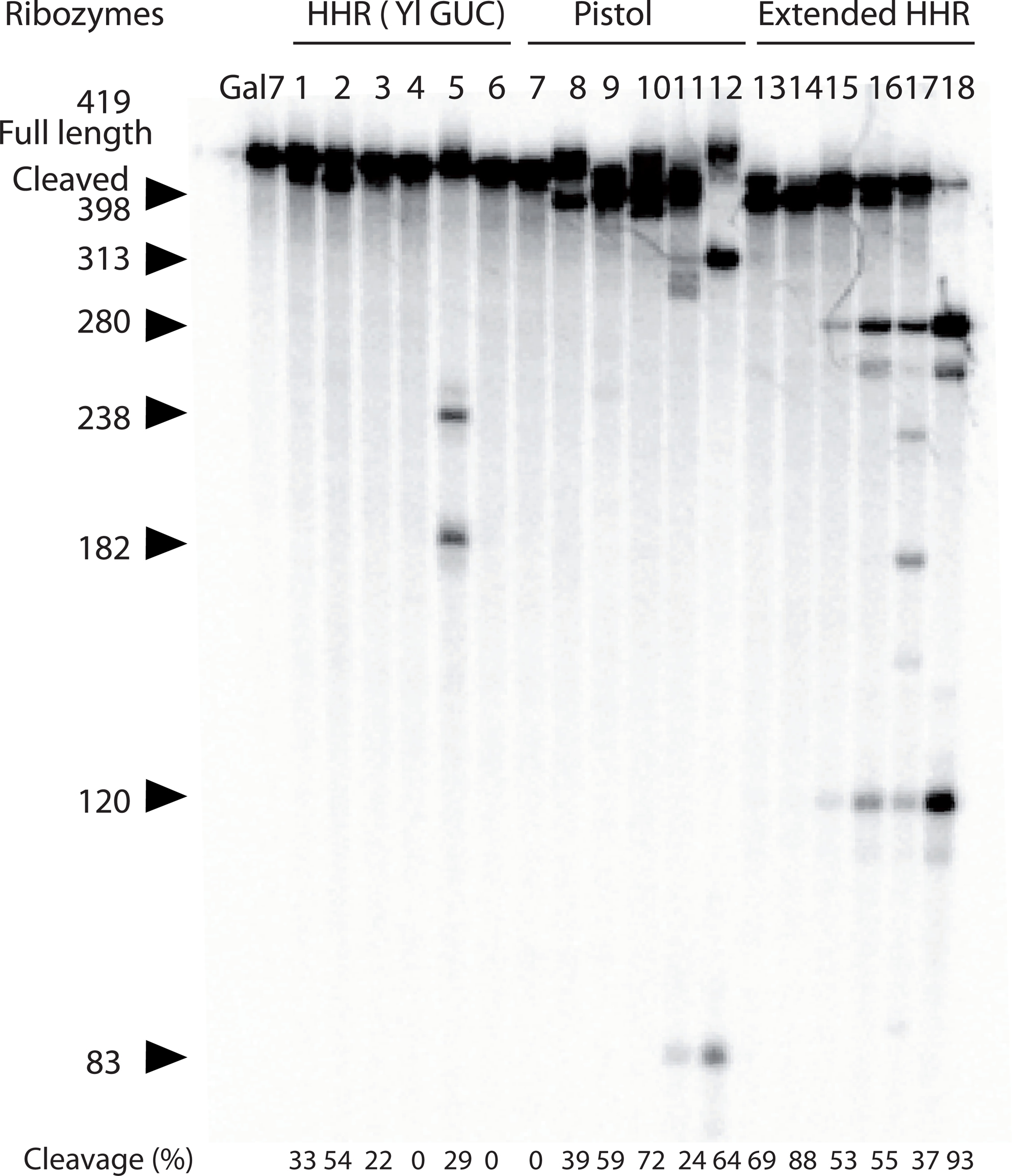
End point cleavage assays of ribozymes against Gal7. Gal7 mRNA labelled with alpha-^32^P-UTP was incubated with each of the eighteen ribozymes designed by Ribosoft (Pistol and HHR ribozymes). The black triangles correspond to the cleaved fragments of each target. The percentages in the bottom of the gel show the cleavage extent of each ribozyme.

### Kinetics measurements

The end-point cleavage assays are good indicators of ribozyme activity, but they are not enough to determine the cleavage rate of a ribozyme. This rate depends on the speed of the reaction, and significant differences can be observed between ribozymes, when comparing them for rate *vs* cleavage extent. In order to assess the rate constant of each active ribozyme, the kinetics were followed for 64 minutes in the same conditions used for the end-point cleavage assays. The kinetics measurements were essentially performed in order to compare each pair of ribozymes for a given parameter evaluated by Ribosoft 2.0 (e.g. “accessibility”) with the highest and the lowest score. This would in principle allow us to figure out if there is a correlation between those parameters and the cleavage efficiency of a ribozyme.

The kinetics were followed for as well (all *k*_obs_ are in table 1). The cleavage rates (*k*_obs_) were as high as 1.8 min^-1^ and most active ribozymes cleaved at > 0.2 min^-1^. Given these results, we could see little to no correlation between ribozyme type and ribozyme activity (Figure 7), or any of the other scores determined by Ribosoft 2.0 because ribozymes with the highest *k*_obs_ are distributed across different types and different accessibility/ temperature/structure scores.

**Figure 7:**
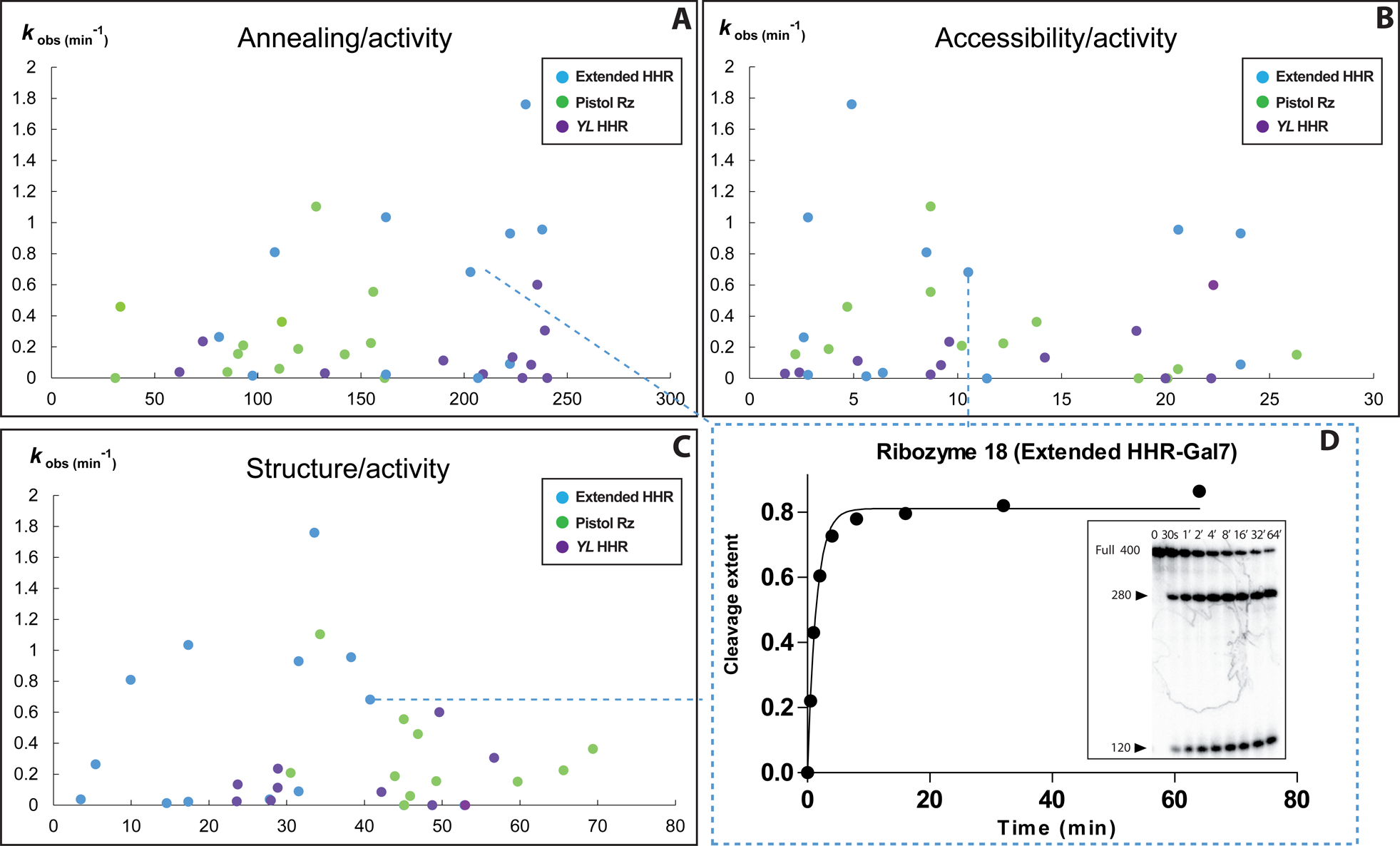
Scatter plots showing the *k*_obs_ of each ribozyme compared to the score of each one of the parameters given by Ribosoft 2.0 : (A) the temperature score relevant for annealing of the ribozyme binding arms to its complementary target, (B) the accessibility score and (C) the structure score. In all three cases, the units of the X axis are arbitrary and correspond to their respective scores. (D) Example of a one phase decay curve (from ribozyme 18) used to calculate the *k*_obs_ of each ribozyme. The inset shows the corresponding gel of ribozyme kinetics used to measure the cleavage extent of the target during a time course over 64 minutes. FMN aptazymes are not represented in that figure.

**Table 1:** Ribozyme activity.

| Rz # | Ribozymes | % | k<sub>obs</sub> | Rz # | Ribozymes | % | k<sub>obs</sub> |
| --- | --- | --- | --- | --- | --- | --- | --- |
| 1 | YL GUC1 (Gal7) | 8 | 0.03796 | 31 | YL GUC1b (Bcl2) | 1 | 0.2349 |
| 2 | YL GUC2 (Gal7) | 48 | 0.08522 | 32 | YL GUC2b (Bcl2) |  | 0 |
| 3 | YL GUC3 (Gal7) | 20 | 0.02993 | 33 | YL GUC3b (Bcl2) | 41 | 0.1121 |
| 4 | YL GUC4 (Gal7) |  | 0 | 34 | YL GUC4b (Bcl2) | 80 | 0.5992 |
| 5 | YL GUC5 (Gal7) | 23 | 0.02336 | 35 | YL GUC5b (Bcl2) | 61 | 0.1331 |
| 6 | YL GUC6 (Gal7) |  | 0 | 36 | YL GUC6b (Bcl2) | 66 | 0.3052 |
| 7 | Pistol1 (Gal7) |  | 0 | 37 | Pistol1b (Bcl2) | 49 | 0.4587 |
| 8 | Pistol2 (Gal7) | 15 | 9.49E-06 | 38 | Pistol2b (Bcl2) | 34 | 0.554 |
| 9 | Pistol3 (Gal7) | 77 | 0.187 | 39 | Pistol3b (Bcl2) | 44 | 0.1539 |
| 10 | Pistol4 (Gal7) | 58 | 0.1509 | 40 | Pistol4b (Bcl2) | 75 | 0.0583 |
| 11 | Pistol5 (Gal7) | 83 | 1.103 | 41 | Pistol5b (Bcl2) | 41 | 0.2084 |
| 12 | Pistol6 (Gal7) | 14 | 0.3625 | 42 | Pistol6b (Bcl2) | 42 | 0.224 |
| 13 | Extend1 (Gal7) | 79 | 0.2644 | 43 | Extend1b (Bcl2) | 30 | 0.01465 |
| 14 | Extend2 (Gal7) | 85 | 1.761 | 44 | Extend2b (Bcl2) | 34 | 0.9564 |
| 15 | Extend3 (Gal7) | 10 | 0.02365 | 45 | Extend3b (Bcl2) | 44 | 1.035 |
| 16 | Extend4 (Gal7) | 6 | 0.09024 | 46 | Extend4b (Bcl2) | 73 | 0.9308 |
| 17 | Extend5 (Gal7) | 4 | 0.03763 | 47 | Extend5b (Bcl2) | 4 | 0.8098 |
| 18 | Extend6 (Gal7) | 86 | 0.6823 | 48 | Extend6b (Bcl2) |  | 0 |
| 19 | FMN act1 (Gal7) | 3 | 0.05396 |  |  |  |  |
| 20 | FMN act2 (Gal7) |  | 0 |  |  |  |  |
| 21 | FMN act3 (Gal7) |  | 0 |  |  |  |  |
| 22 | FMN act4 (Gal7) |  | 0 |  |  |  |  |
| 23 | FMN act5 (Gal7) |  | 0 |  |  |  |  |
| 24 | FMN act6 (Gal7) | 25 | 0.009182 |  |  |  |  |
| 25 | FMN inh1 (Gal7) |  | 0 |  |  |  |  |
| 26 | FMN inh2 (Gal7) | 9 | 1.34E-05 |  |  |  |  |
| 27 | FMN inh3 (Gal7) | 2 | 0.01282 |  |  |  |  |
| 28 | FMN inh4 (Gal7) | 15 | 6.14E-06 |  |  |  |  |
| 29 | FMN inh5 (Gal7) | 4 | 0.003411 |  |  |  |  |
| 30 | FMN inh6 (Gal7) |  | 0.00 |  |  |  |  |

%= percentage cleaved after 64 min of incubation at 37°C

### FMN Aptazymes

For FMN activated and FMN inhibited aptazymes, twelve ribozymes targeting Gal7 mRNA were tested in the presence and absence of 200 μM of FMN. For the FMN-activated version of the aptazyme, two of the six ribozymes were active under both conditions (presence and absence of FMN), with cleavage efficiencies of 22 and 57%, while four were inactive. As for the six FMN-inhibited ribozymes, four of the six ribozymes tested were active in the absence of FMN (giving cleavage efficiencies of 10, 13, 38 and 58%), while their activity was completely suppressed when FMN was added to the reaction (Figure 8). Although only four ribozymes have shown the desired activity in presence/absence of FMN, the result still indicates that Ribosoft 2.0 was not only able to design different *trans*-acting ribozymes, but for the first time, design inducible *trans*-acting ribozymes as well.

**Figure 8:**
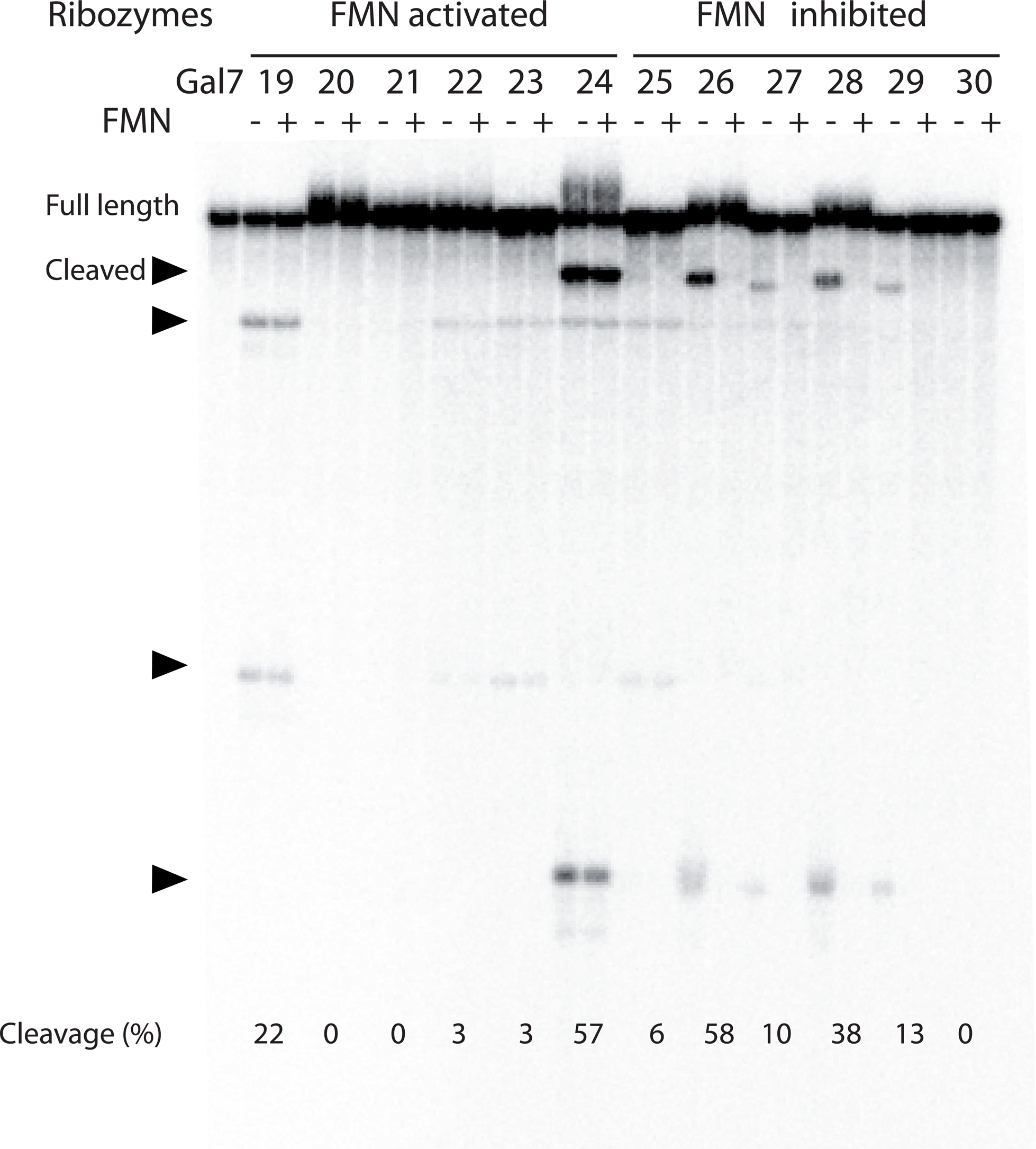
End point cleavage tests of activated and inhibited version of trans acting FMN aptazymes against Gal7 mRNA in presence and absence of FMN. The activated version of the aptazyme did not show any activity except for aptazyme 19 that is active in presence and absence of FMN. The inhibited version : four aptazymes (26, 27, 28 and 29) have shown the desired activity where they were active in absence of FMN and the lost completely their activity when FMN was present in the reaction solution.

## Discussion

Ribosoft 2.0 is a web service with a user-friendly interface offering an easy and simple way to design specific types of ribozymes and, if desired, allows the entry of advanced parameter values. Our results show that the service is an efficient tool for the generation of different types of active ribozymes and aptazymes targeting any desired RNA in an automated fashion, quickly and correctly. This renders Ribosoft 2.0 as the only publicly available web service with such facility. There is no other bioinformatics tool allowing for the generation of this number of different structures simultaneously. To our knowledge, the only tool handling more than one type of ribozyme is ‘Ribosubstrate’ (42) – but it does not automatically design ribozymes. Rather, it requires prior determination of the target site and ribozyme sequence and then uses BLAST to look for off-targets of SOFA-HDV, hammerhead ribozymes or siRNAs. Ribosoft 2.0 has advantages over ‘Aladdin’ (43), a bioinformatics tool that generates solely hammerhead ribozyme designs, relaying on the target sequence as the only input. Moreover, the only parameter calculated with this tool is the annealing temperature (T_m_) of right and left arms of the ribozyme; it does not check for off-targets. In this case, each of the mentioned programs misses an important part of the design, either the specificity parameter or the automation of the design. In our case, we developed a tool combining the options given by both programs, and with advantageous extensions. Many of these extensions were already provided by the first version of Ribosoft (25) that we used to generate HHR against the mRNA of PABPN1, which were active *in vitro,* in mammalian cells and in *C. elegans* (44).

In the new version of the web service as well as the old one, the design is made, taking in account the specificity of a ribozyme by blasting sequences in NCBI. In addition, Ribosoft 2.0 computes three other parameter. The accessibility of the cleavage sites is calculated for all sites compatible with the sequence constraints provided by the ribozyme model. The program uses the secondary structure of the target to check for single stranded regions. It also checks for the quality of the newly designed ribozyme structure and, lastly, it provides a score reflecting the annealing temperature between the target and the ribozyme’s arms.

Regarding a relationship between scores calculated by Ribosoft 2.0 and ribozyme activity, since the software already screens out all ribozymes likely to be inactive, it excludes ribozymes from a wide range of parameter values and predicted efficiency, hindering our attempt to evaluate such a correlation. Instead, we proved that most ribozymes designed by Ribosoft 2.0 (except aptazymes) were active, i.e. the program has generated 8-9 active ribozymes out of every 10 designs, independently of parameter values. Actually, all of the chosen ribozymes were ranked #1 by Ribosoft, so this ranking was not particularly useful. This can be explained by the filters of Ribosoft 2.0, which already eliminate all the ribozymes having minor defaults and lower scores for ribozymes. In the end, this sorts through sequences to select only the highly active ribozymes, even those with the lowest scores.

The main addition to the Ribosoft 2.0 software is the possibility of adding almost any type of ribozyme. It is even possible to include the presence of a pseudoknot within the “ribozyme strand”, although folding of pseudoknots is not formally computed. As for pseudoknots between the “substrate” and “ribozyme” strands, such as for the validated HHRs from *Y. lipolytica*, they are taken into account via the hybridization of each binding arm. To our knowledge, Ribosoft 2.0 allows for the first time to automatically design allosteric ribozymes against desired targets. Indeed, we generated the first *trans*-acting aptazymes and demonstrated activity of the FMN-inhibited HHR aptazymes in presence of FMN. Known examples of *trans*-acting aptazymes act in the following way: the output of the cleavage reaction works as the effector or an input of another reaction. As an example, a guanine aptazyme that generates a pri-miRNA as an output acts in combination with the RISC complex to degrade a target mRNA (45). In our case, the aptazyme works directly as a degradation machine of the target RNA in a sequence specific manner. This is a novelty provided by our software giving the users the opportunity to have on the same RNA molecule a detector of a specific molecule and an effector against a specific target sequence, which we can easily modulate depending on the target. This option will not be limited only to aptazymes, but it can also work for other types of allosteric ribozymes controlled by the presence of a certain input like an RNA or DNA sequence. Despite the fact that our results show only four active aptazymes corresponding to the given condition, they still indicate that for a few easily designed aptazymes users are very likely to get design sequences fitting their needs. Indeed, four out of six of the FMN-inhibited aptazymes against Gal7 mRNA function as expected. As for the FMN-activated aptazymes, the fact that none of them worked may indicate that the structure model described by Soukup and Breaker (46) does not correspond exactly to the active structure of the aptazyme, which would lead Ribosoft 2.0 to inappropriately score the structure of the ribozyme. These RNAs can be used in many applications like the replacement of the *cis*-acting aptazymes included upstream or downstream of the target gene in many previous examples of aptazyme use in synthetic biology (19,47–49). By having *trans*-acting aptazymes, the function of aptazymes can be applied to prokaryotic and eukaryotic cells, including mammalian and human cells, without the need of combining different macromolecules (*cis*-acting aptazyme and RISC complex with miRNA or siRNA).

The results we got from aptazymes were limited because all of the aptazymes against Blc2 mRNA were inactive (data not shown). Also some aptazymes against Gal7 mRNA were active but did not show any difference in activity in the presence or absence of FMN. A possible explanation is that these aptazymes were developed using minimal HHRs which are 100 fold less active than full-length HHRs. Besides, the input secondary structure given to Ribosoft 2.0 for the design of these aptazymes is only representative of either the active or the inactive state of the aptazyme in presence or absence of a specific ligand. The structure for the other state might be important missing information that Ribosoft 2.0 needs, to make the comparison between the two states, and design ribozymes with properly optimized structures. In that regard, the native conventional HHR is the most studied catalytic RNA and it has a well characterized structure, contrary to aptazymes which are built based on aptamers selected *in vitro* and usually having poorly characterized structures. In other words, the biggest limitation of Ribosoft 2.0 for the design of aptazymes might not be the algorithm underlying the software, but rather the lack of comprehensive characterization of most selected aptamers and aptazymes. This can be circumvented by using well-studied examples, by characterizing aptamers/aptazymes that bind ligands of interest, or simply by including information on secondary structure only for the “ribozyme portion” and leaving the aptamer portion as if it was single-stranded. Nevertheless, the published FMN aptazyme structures allowed us to demonstrate the applicability of Ribosoft 2.0 for aptazymes. In the future, any aptazyme or other allosteric ribozymes with a known structure can be used as input to Ribosoft 2.0, as the software allows for future addition to new ribozyme templates.

## Acknowledgements

The authors wish to thank Atef Nehdi for helpful discussion.

## Supplementary material

### Ribozyme models

#### *Yarrowia lipolytica* hammerhead ribozyme (*cis* version has a pseudoknot)

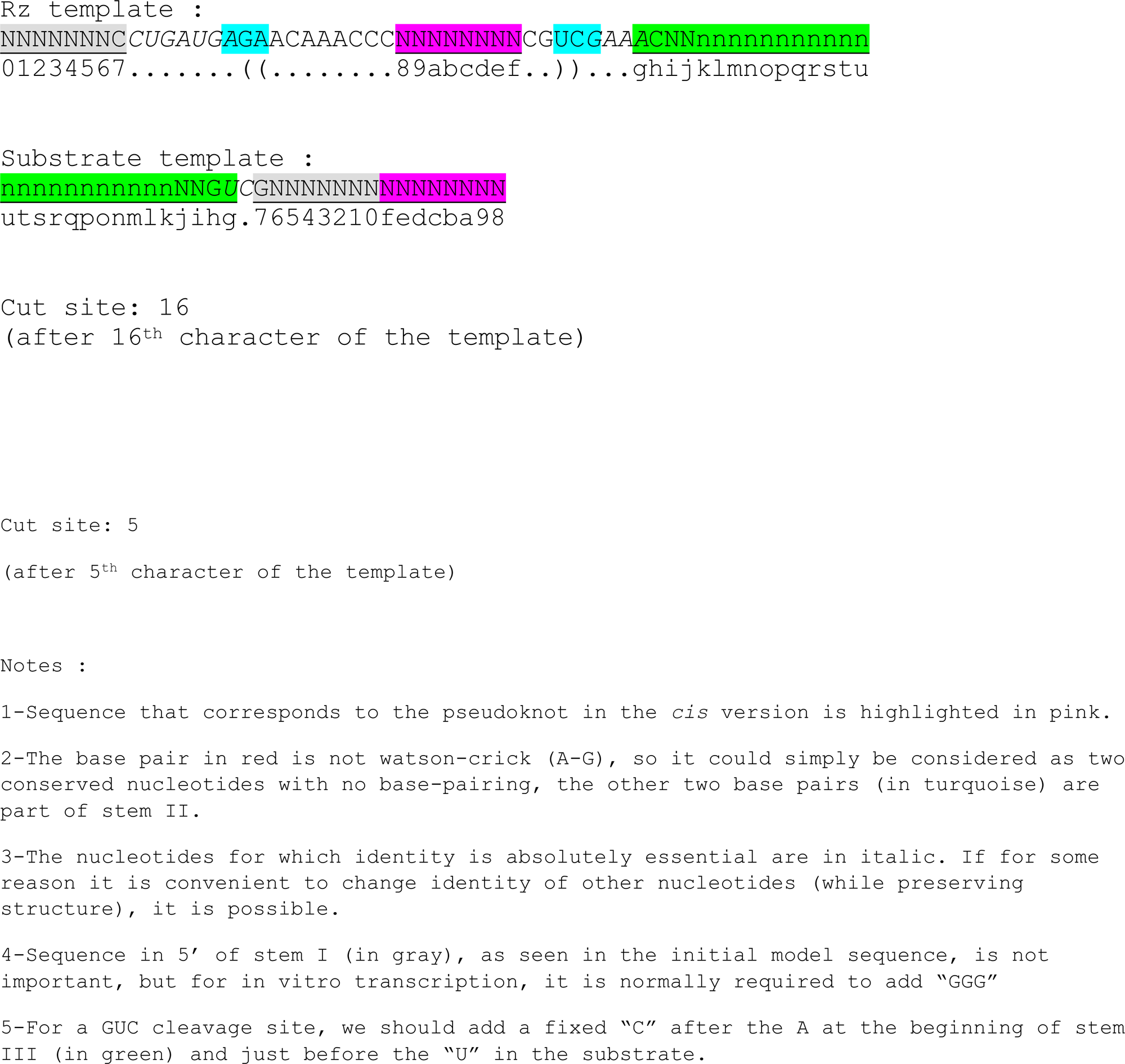

#### Extended Hammerhead ribozyme

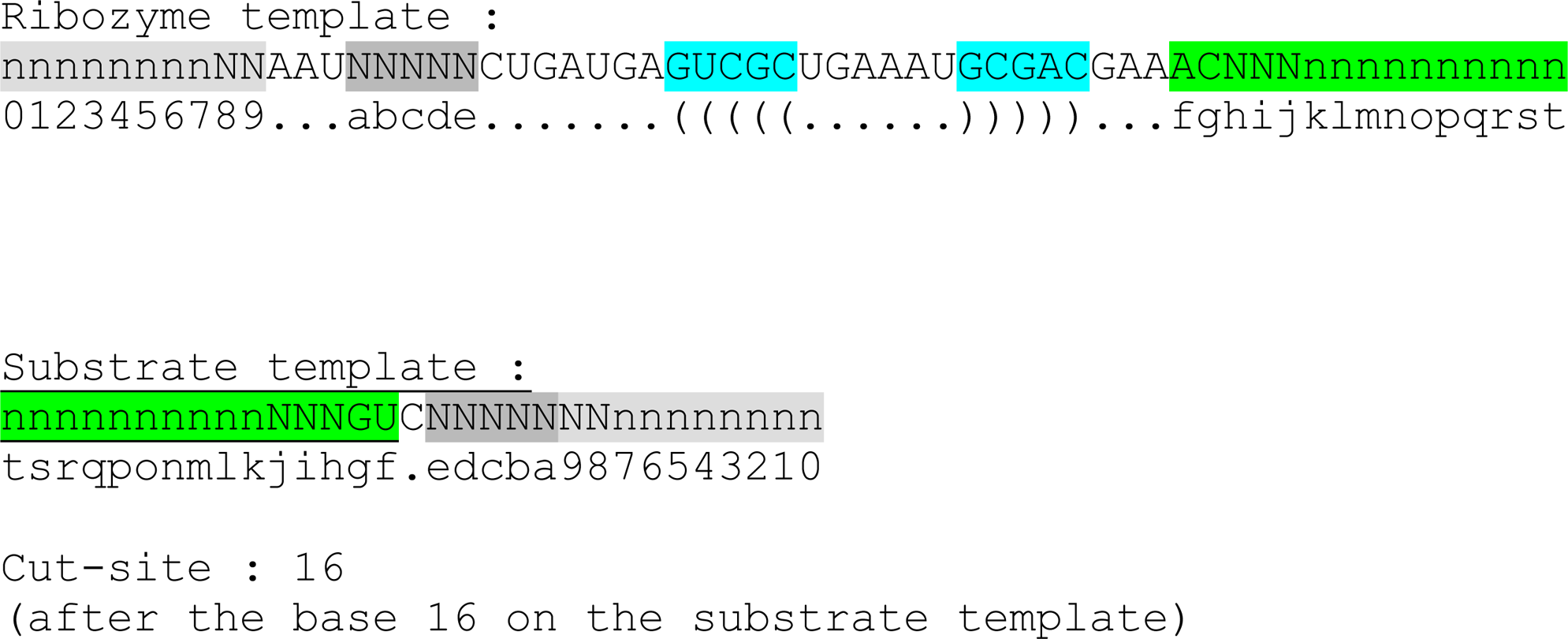

#### Pistol ribozyme

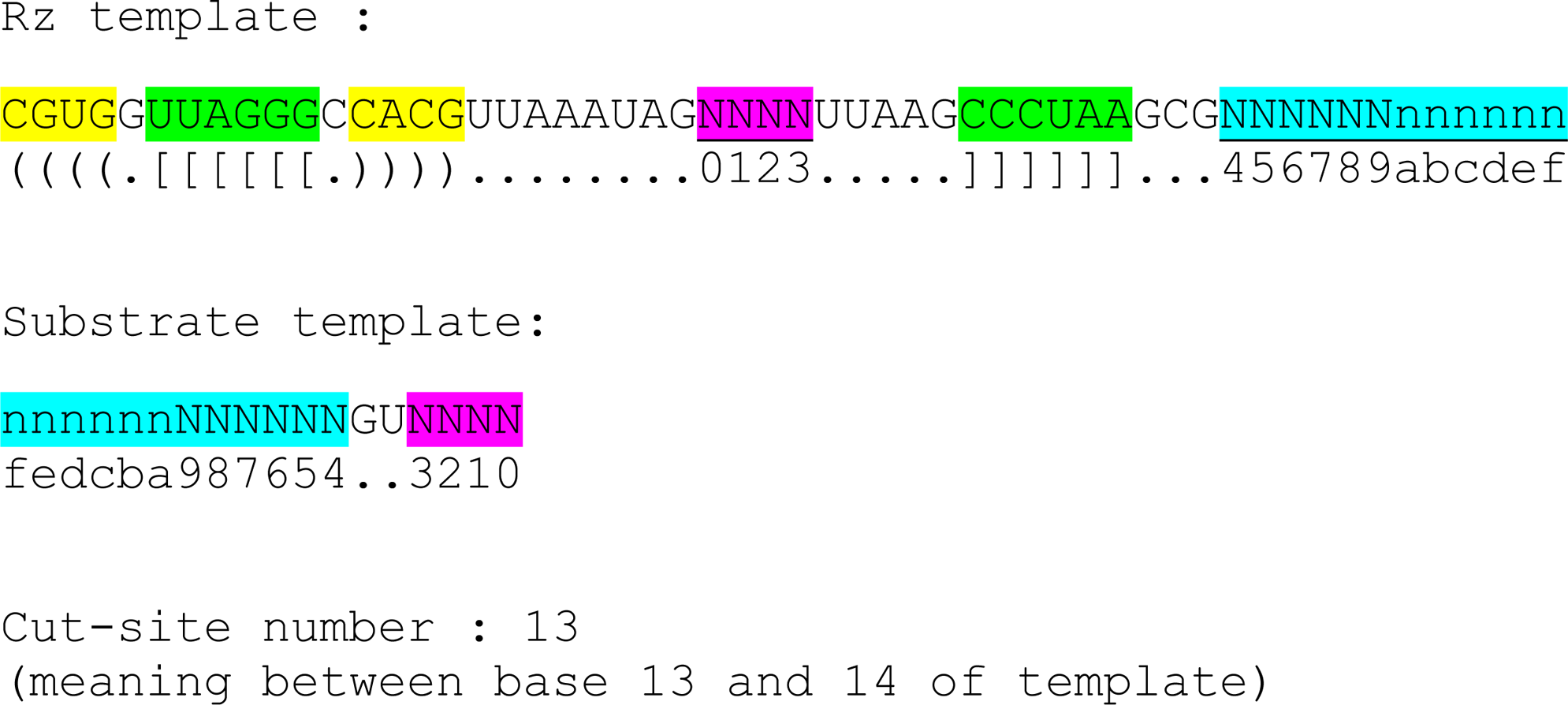

#### Hammerhead induced by FMN

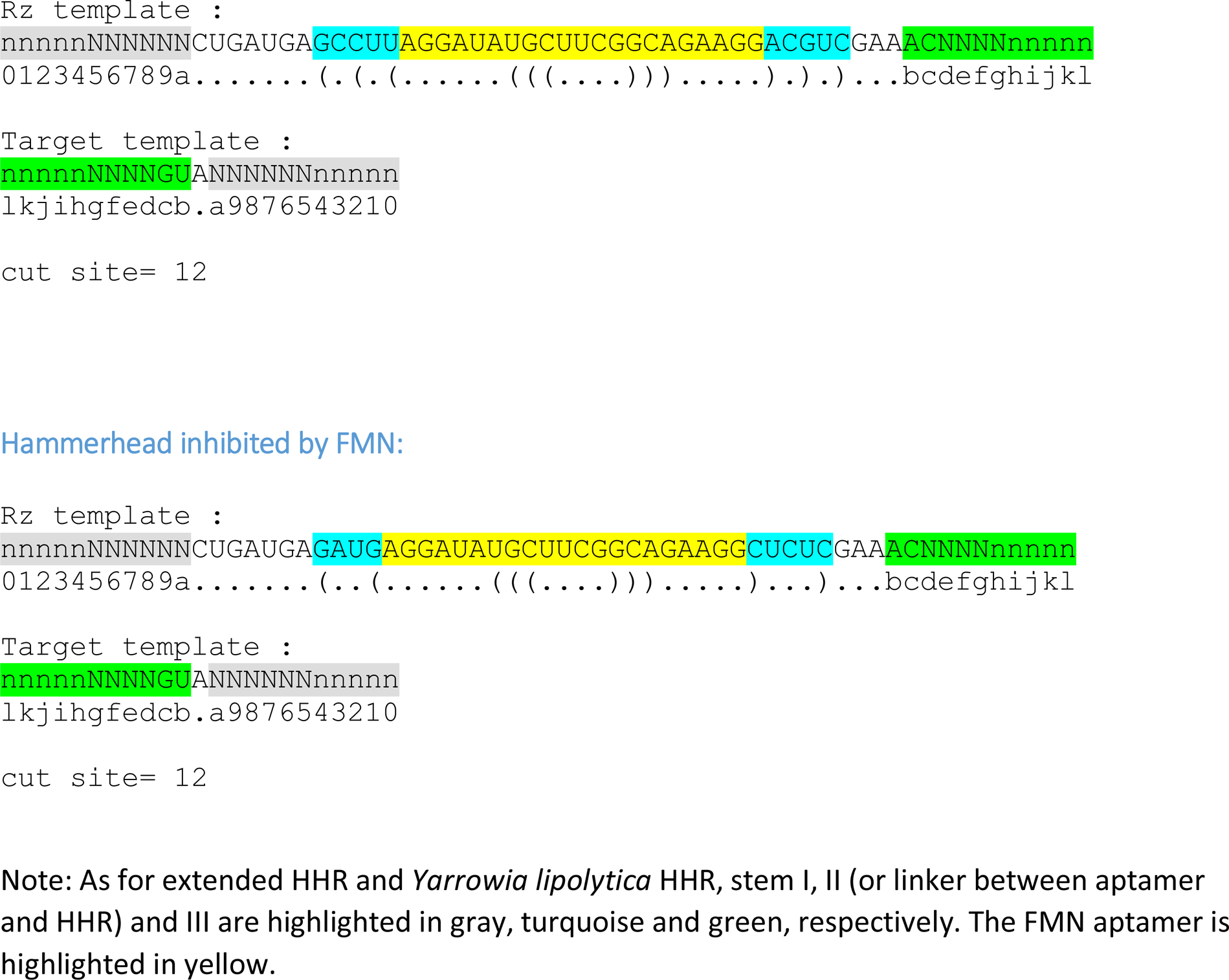

#### Hammerhead inhibited by FMN

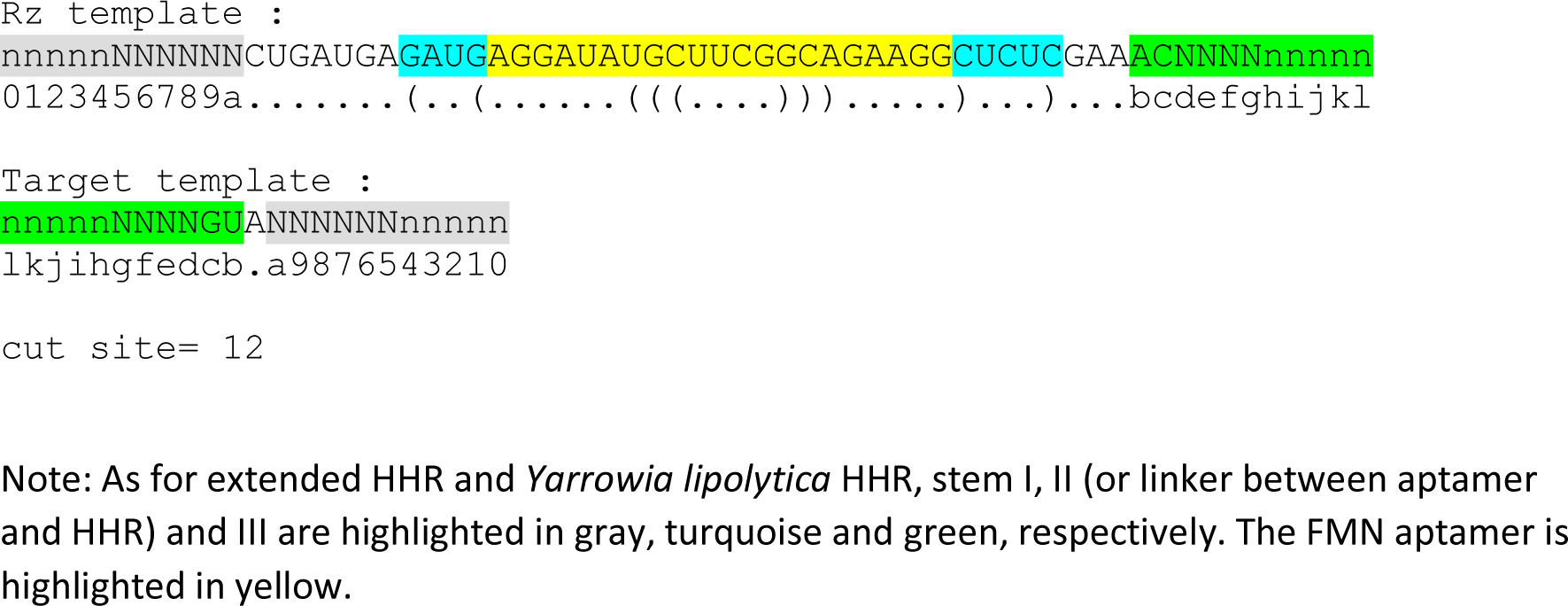

**Table S1:**
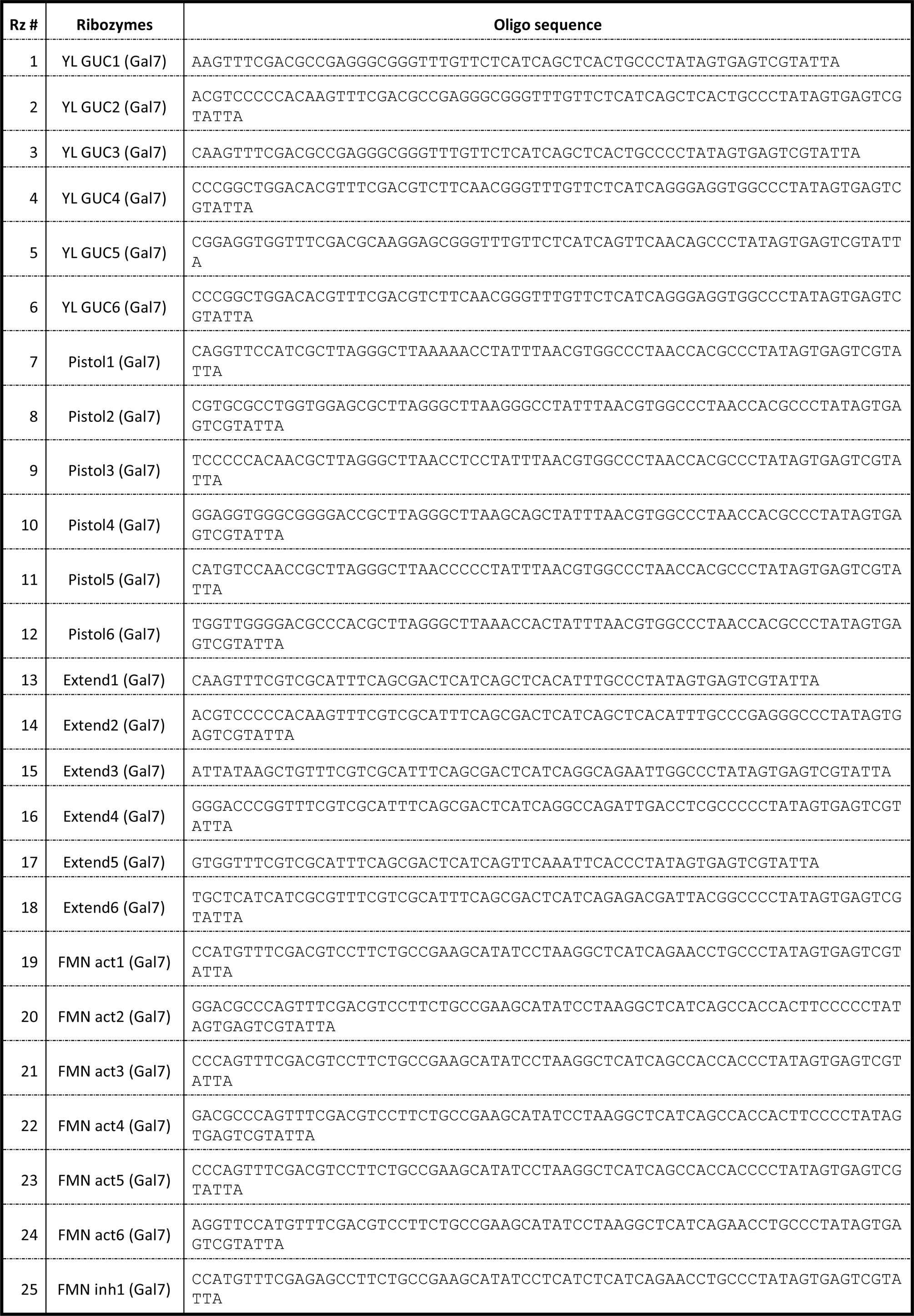
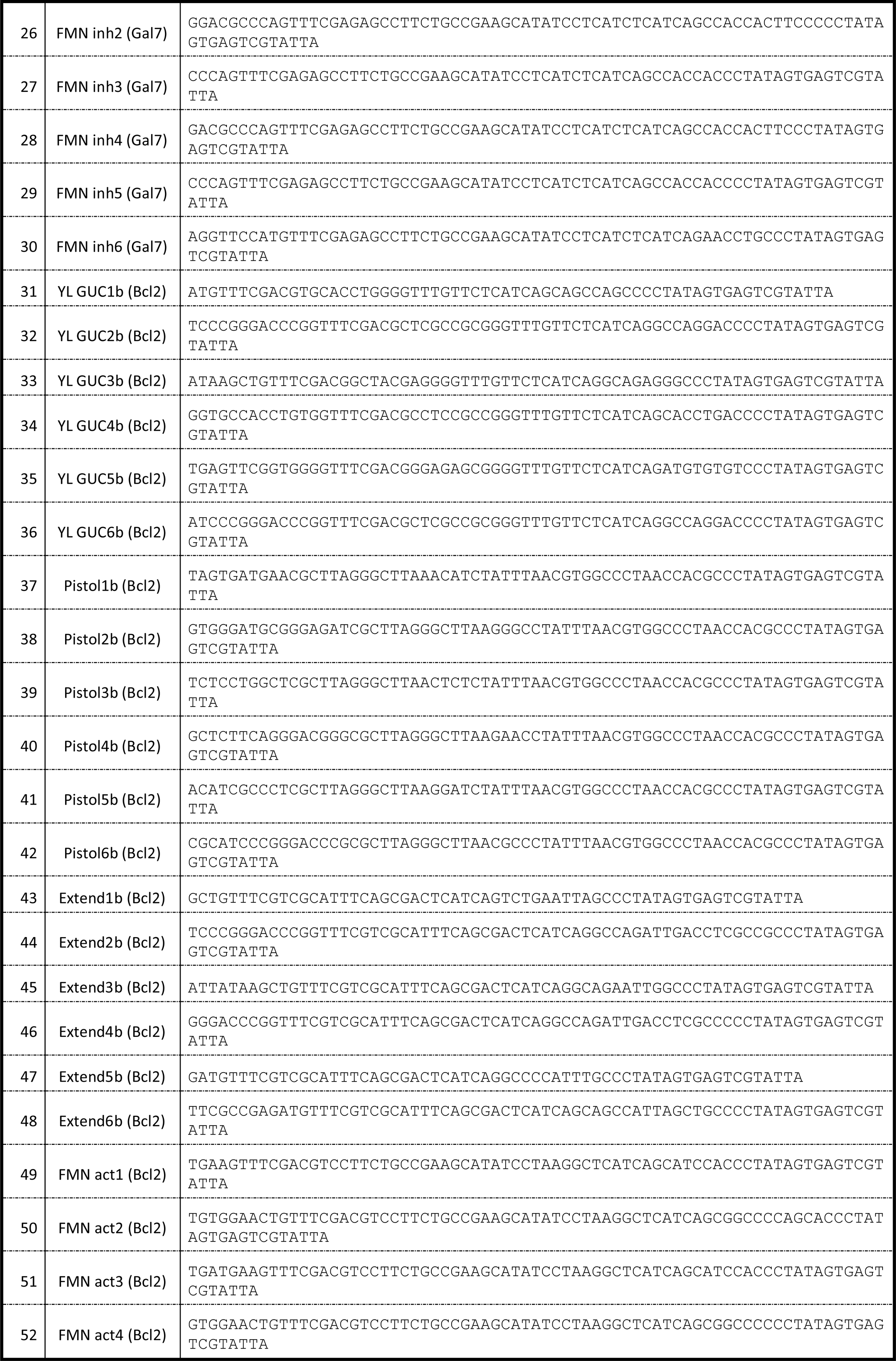
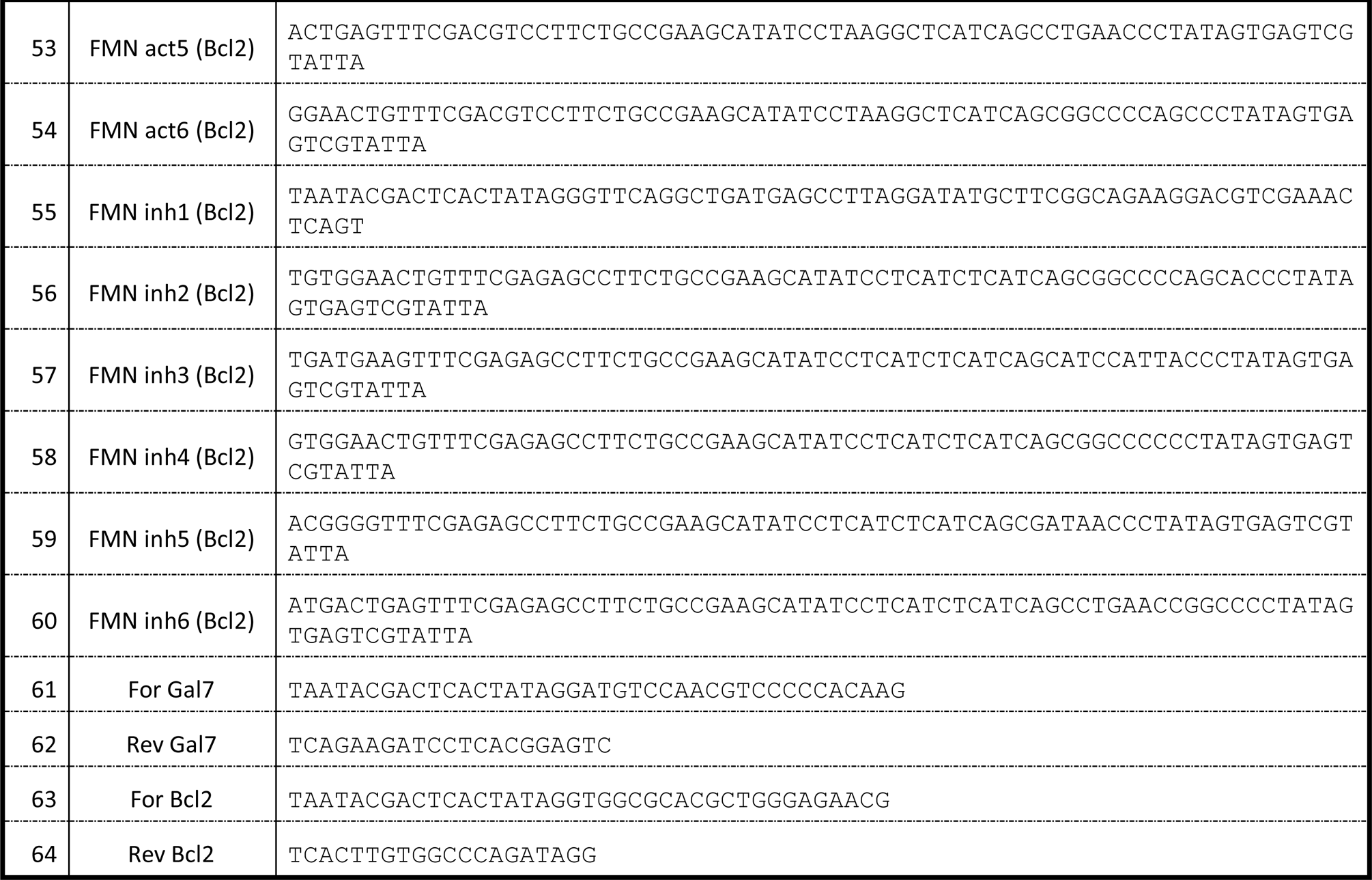
Oligonucleotides sequences.

**Table S2:** Parameters scores of chosen ribozymes.

| Rz # | Ribozymes | Parameters | Score |
| --- | --- | --- | --- |
| 1 | YL GUC1 (Gal7) | Tm min | 62.055 |
| 2 | YL GUC2 (Gal7) | Tm max | 232.478 |
| 3 | YL GUC3 (Gal7) | Accessibility min | 1.7 |
| 4 | YL GUC4 (Gal7) | Accessibility max | 22.2 |
| 5 | YL GUC5 (Gal7) | Structure min | 23.593 |
| 6 | YL GUC6 (Gal7) | Structure max | 48.722 |
| 7 | Pistol1 (Gal7) | Tm min | 30.933 |
| 8 | Pistol2 (Gal7) | Tm max | 161.474 |
| 9 | Pistol3 (Gal7) | Accessibility min | 3.8 |
| 10 | Pistol4 (Gal7) | Accessibility max | 26.3 |
| 11 | Pistol5 (Gal7) | Structure min | 34.292 |
| 12 | Pistol6 (Gal7) | Structure max | 69.411 |
| 13 | Extend1 (Gal7) | Tm min | 81.307 |
| 14 | Extend2 (Gal7) | Tm max | 229.866 |
| 15 | Extend3 (Gal7) | Accessibility min | 2.8 |
| 16 | Extend4 (Gal7) | Accessibility max | 23.6 |
| 17 | Extend5 (Gal7) | Structure min | 3.525 |
| 18 | Extend6 (Gal7) | Structure max | 40.736 |
| 19 | FMN act1 (Gal7) | Tm min | 98.405 |
| 20 | FMN act2 (Gal7) | Tm max | 207.06 |
| 21 | FMN act3 (Gal7) | Accessibility min | 8.4 |
| 22 | FMN act4 (Gal7) | Accessibility max | 22.4 |
| 23 | FMN act5 (Gal7) | Structure min | 29.651 |
| 24 | FMN act6 (Gal7) | Structure max | 52.461 |
| 25 | FMN inh1 (Gal7) | Tm min | 98.405 |
| 26 | FMN inh2 (Gal7) | Tm max | 207.06 |
| 27 | FMN inh3 (Gal7) | Accessibility min | 8.4 |
| 28 | FMN inh4 (Gal7) | Accessibility max | 22.4 |
| 29 | FMN inh5 (Gal7) | Structure min | 29.478 |
| 30 | FMN inh6 (Gal7) | Structure max | 51.639 |
| 31 | YL GUC1b (Bcl2) | Tm min | 73.379 |
| 32 | YL GUC2b (Bcl2) | Tm max | 240.264 |
| 33 | YL GUC3b (Bcl2) | Accessibility min | 5.2 |
| 34 | YL GUC4b (Bcl2) | Accessibility max | 22.3 |
| 35 | YL GUC5b (Bcl2) | Structure min | 23.711 |
| 36 | YL GUC6b (Bcl2) | Structure max | 56.681 |
| 37 | Pistol1b (Bcl2) | Tm min | 33.491 |
| 38 | Pistol2b (Bcl2) | Tm max | 155.947 |
| 39 | Pistol3b (Bcl2) | Accessibility min | 2.2 |
| 40 | Pistol4b (Bcl2) | Accessibility max | 20.6 |
| 41 | Pistol5b (Bcl2) | Structure min | 30.511 |
| 42 | Pistol6b (Bcl2) | Structure max | 65.625 |
| 43 | Extend1b (Bcl2) | Tm min | 97.492 |
| 44 | Extend2b (Bcl2) | Tm max | 237.865 |
| 45 | Extend3b (Bcl2) | Accessibility min | 2.8 |
| 46 | Extend4b (Bcl2) | Accessibility max | 23.6 |
| 47 | Extend5b (Bcl2) | Structure min | 9.931 |
| 48 | Extend6b (Bcl2) | Structure max | 45.15 |
| 49 | FMN act1 (Bcl2) | Tm min | 85.063 |
| 50 | FMN act2 (Bcl2) | Tm max | 204.685 |
| 51 | FMN act3 (Bcl2) | Accessibility min | 3.3 |
| 52 | FMN act4 (Bcl2) | Accessibility max | 14.6 |
| 53 | FMN act5 (Bcl2) | Structure min | 31.579 |
| 54 | FMN act6 (Bcl2) | Structure max | 55.275 |
| 55 | FMN inh1 (Bcl2) | Tm min | 85.063 |
| 56 | FMN inh2 (Bcl2) | Tm max | 204.685 |
| 57 | FMN inh3 (Bcl2) | Accessibility min | 3.3 |
| 58 | FMN inh4 (Bcl2) | Accessibility max | 14.6 |
| 59 | FMN inh5 (Bcl2) | Structure min | 26.714 |
| 60 | FMN inh6 (Bcl2) | Structure max | 51.762 |

Ribozymes are listed by pairs of high/low values for the indicated parameter computed by Ribosoft 2.0.

